# Decoding state specific connectivity during speech production and perception

**DOI:** 10.1101/2025.10.10.681678

**Authors:** Yasamin Esmaeili, Amirhossein Khalilian-Gourtani, Orrin Devinsky, Werner K. Doyle, Patricia Dugan, Daniel Friedman, Adeen Flinker

**Author notes:** **For correspondence:** (YE); (AF).

## Abstract

Understanding how dynamic brain networks support language perception and production is central to cognitive neuroscience. A vast network-based literature has employed functional connectivity (FC), primarily using resting-state and task-based fMRI. However, methodological limitations have hindered this approach in language processing, particularly during speech production. Here, we address this gap by employing a large cohort of electrocorticographic (ECoG) patients (N = 38) to investigate the networks driving speech perception and production. We acquired data while patients were engaged in a controlled battery of speech production tasks focusing on five behavioral states (auditory perception, picture perception, reading perception, speech production, and baseline). Using linear classifiers we were able to robustly decode behavioral states from single-trial FC (i.e. Pearson correlations) of the neural activity patterns, achieving a mean accuracy of 73.9%. These classifiers revealed distinct network signatures underlying auditory and visual perception as well as speech production via stable network connectivity. Importantly, the network signatures included both regions with robust local neural activity and those with minimal or no detectable activation. Such signatures indicate that even low-activity regions contribute critically to differentiating behavioral states. Further, our findings implicate connectivity between anterior temporal lobe and perisylivan sites as signatures driving speech production classification. Together, these results underscore the significance of functional connectivity analysis as a complementary dimension to investigating local neural activity, and suggest that the functional networks supporting speech extend beyond the classical, most metabolically active regions.

## Introduction

Human cognition and behavior is coordinated by a complex network of interconnected brain regions (***Sporns and Betzel, 2016***; ***Bassett and Sporns, 2017***). A large body of literature has described these networks leveraging functional connectivity (FC) across regions during rest (***Biswal et al., 1995***; ***Fox et al., 2005***; ***Greicius et al., 2009***) as well as cognitive processing (***Friston et al., 1993***; ***Cole et al., 2014***, ***2021***). Resting state data has identified reproducible intrinsic networks of functionally connected regions, such as the default mode, dorsal attention, and sensorimotor network, that resemble FC patterns observed during tasks (***Biswal et al., 1995***; ***Fox and Raichle, 2007***; ***Smith et al., 2009***). While task-based FC studies are able to reproduce intrinsic network, they also provide evidence for network reconfiguration during changing cognitive demand (***Cole et al., 2014***; ***Shine et al., 2016***; ***Hearne et al., 2017***).

One hallmark of human behavior is our ability to produce complex and rapid speech utterances. However, there are only a select few studies examining functional connectivity during the production of speech and language (***Fuertinger et al., 2015***; ***Riecker et al., 2005***; ***Geranmayeh et al., 2014***; ***Ewald et al., 2012***). This is mainly due to the severe motor artifacts introduced by the jaw, tongue, and laryngeal movements during speech that are inherent to noninvasive neuroimaging and electrophysiological recordings (***Grabski et al., 2012***; ***Abbasi et al., 2021***). As a result, the network principles that support the rapid sensorimotor transformations of speech are still poorly understood. However, human invasive electrocorticographic (ECoG) recordings circumvent these motor artifacts. Intracranial signals are obtained during neurosurgical procedures directly from the depth and surface of the brain providing a high spatial resolution and millisecond temporal precision. The vast majority of previous intracranial studies have focused on local high-gamma neural activity as a proxy for population firing (***Mesgarani et al., 2014***; ***Pasley et al., 2012***; ***Edwards et al., 2010***; ***Flinker et al., 2015***; ***Chang et al., 2010***; ***Bouchard et al., 2013***; ***Mugler et al., 2014***; ***Cheung et al., 2016***). These studies have profoundly advanced our understanding of how local neural activity encodes articulatory gestures and phonetic features (***Bouchard et al., 2013***; ***Mesgarani et al., 2014***; ***Chang et al., 2010***; ***Mugler et al., 2014***; ***Cheung et al., 2016***), and how speech production sequences involve distinct temporal activation patterns across language-relevant cortical regions (***Edwards et al., 2010***; ***Flinker et al., 2015***). However, the dynamics of inter-regional connectivity during speech production tasks remain largely unexplored.

Measuring functional connectivity in ECoG typically involves assessing synchronous cofluctuations in broadband amplitude, or within specific frequency bands, most often high-gamma (i.e. 70-150 Hz frequency band), using pairwise correlation or coherence (***Axmacher et al., 2010***; ***Kucyi et al., 2018***). Analyses of long-duration resting-state recordings have revealed stable correlation templates that closely mirror canonical functional Magnetic Resonance Imaging (fMRI) networks (***Kramer et al., 2011***; ***Kucyi et al., 2018***). During overt speech production, high-gamma coherence between inferior frontal gyrus and superior temporal cortex increases selectively during active vocalization relative to passive listening (***Kingyon et al., 2015***), underscoring the value of correlation-based FC for probing rapid sensorimotor integration. Yet most FC studies in ECoG still rely on long analysis windows or condition-averaged data. This practice can mask the transient network dynamics that support rapid behaviors such as speech. Undirected FC has also been rarely applied to single-trial decoding of behavioral stages, so low-amplitude but behaviorally relevant connectivity patterns are often lost in averaging. As a result, we still do not understand which inter-regional connectivity patterns are specific to different successive stages of speech perception and production, or whether these interactions reveal network structure invisible to analyses that focus only on local activity.

Here, we directly address these gaps by examining single-trial functional connectivity dynamics in ECoG recordings from a cohort of 38 participants performing speech-production tasks. We employ a robust machine learning approach—support vector machine (SVM) classification—to decode behavioral states from trial-specific FC patterns, and we translate the resulting weights into interpretable connectivity signatures using forward model transformations (***Haufe et al., 2014***). We aim to address two main questions: (i) Are the cortical networks supporting speech perception and production decodable based on connectivity patterns alone? And (ii) are the associated discriminative connections specific to regions with high neural activity, or do they include network structures obscured by analyses focusing solely on activity?

## Results

### Single-Trial Functional Connectivity

To investigate the dynamics of single trial functional connectivity during speech perception and production, we collected electrocorticography (ECoG) data, which provide high spatiotemporal resolution. We recorded neural activity from 38 participants while performing three word production tasks (see Experimental Tasks and Behavioral States, ***Figure 1***A and B) and defined five behavioral states of interest (***Figure 1***C): Auditory perception, picture perception, reading perception, speech production, and baseline (production and baseline are shared across tasks).

**Figure 1.**
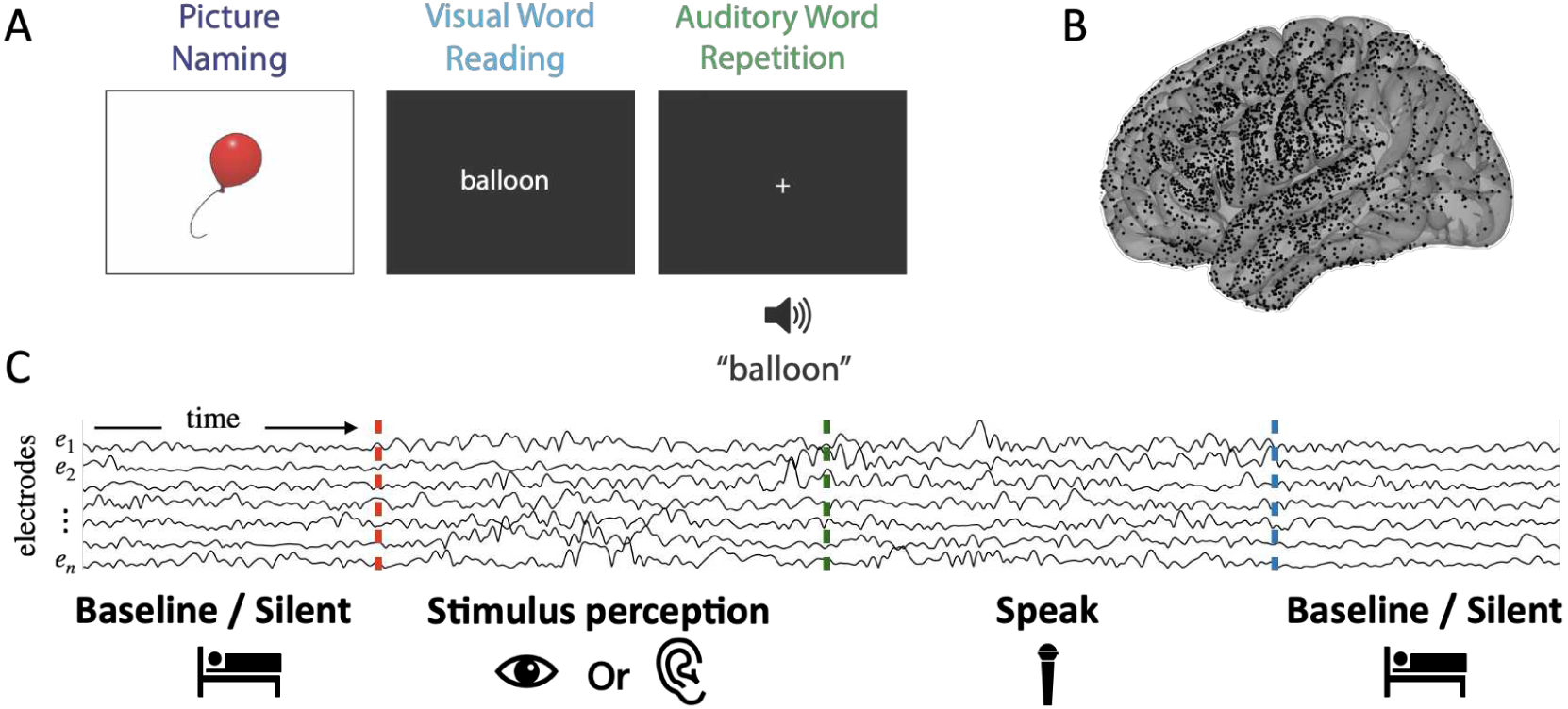
Overview of task and data structure. (A) Battery of language tasks that all culminate in the same speech utterance. The full list of stimuli and associated number of phonemes and syllables are included in **Supplementary Table 1**. (B) Electrode coverage across participants; electrodes projected on the left hemisphere (separate right and left hemisphere coverage is included in **Supplementary Figure 1**) (C) The sequence of the behavioral states of interest in a single trial.

Analyzing functional connectivity (FC) at the single-trial level can reveal nuanced dynamics, including weak yet consistent interactions, that may be obscured during the common practice of trial-averaging (***He et al., 2008***; ***Solomon et al., 2018***; ***Huang et al., 2023***). We segmented all single-trial neural activity into state-specific intervals and computed FC for each (see Single-Trial Functional Connectivity). In order to provide a view of single-trial connectivity, we create “connectivity vectors” comprised of the upper-triangular elements of the FC matrices (***Figure 2***A, left). The entries of these vectors represent all connections between pairs of cortical electrodes. Grouping the connectivity vectors by behavioral state provides a comprehensive view of the connectivity patterns. We illustrate this for one participant, revealing substantial trial-to-trial variability (***Figure 2***A, right). Some states (e.g. picture and reading perception) show obviously similar frontal–frontal and occipital–frontal connections. However, we were interested in identifying patterns of connectivity that are distinct across states.

**Figure 2.**
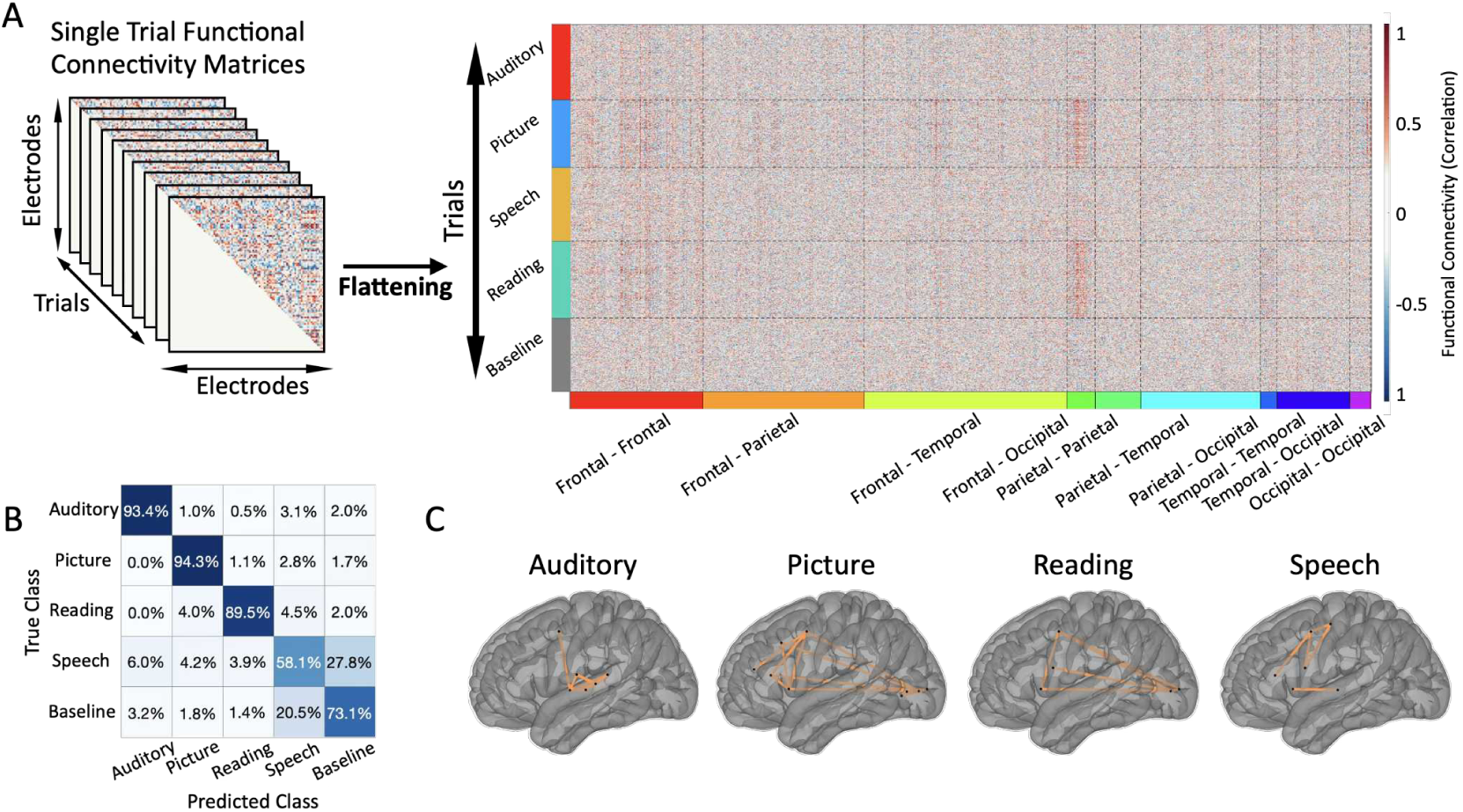
Functional Connectivity analysis for an exemplar participant. (A) Demonstration of flow of creating the functional connectivity feature and resulting vectors across all states. The values are ordered based on connections between different brain lobes (e.g. Frontal-Frontal, Frontal-Parietal, etc.). The features show clear differences in the connectivity patterns of the behavioral states. (B) Confusion matrix showing high accuracy classification of the connectivity vectors. This matrix was calculated using 5-fold cross-validation on the model for the exemplar participant. (C) Discriminative connections for each behavioral state. Orange lines depict connections whose median absolute deviation of the transformed SVM weights survive a Laplacian tail bound significance threshold; node positions correspond to electrodes.

In order to test whether these patterns contained sufficient information to differentiate behavioral states we employed a decoding approach. We trained a multi-class Support Vector Machine (SVM) classifier on the connectivity vectors in a representative participant, achieving a mean cross-validated accuracy of 79.02% (chance level is 20%, ***Figure 2***B). The performance of the model provides evidence for robust state-specific connectivity patterns in this participant. In order to identify these state-specific connections we examined the model weights. We assessed significance of weights after following common practices of transformation (***Haufe et al., 2014***), to ensure unbiased interpretability (see Decoding Framework). This identified specific networks driving classification performance, for example intra-auditory-cortex connections during auditory perception (***Figure 2***C). We call these *discriminative connections*, providing a distinction for connectivity patterns that significantly differentiate behavioral states.

### State-Specific Connectivity Patterns Across Participants

To test whether the state-specific connectivity patterns observed in the representative participant generalize to the entire cohort, we replicated our decoding pipeline within each of the N = 38 participants. Across all participants, the classifiers performed robustly (***Figure 3***A): perceptual states were decoded with a mean of 88.7-90.9% accuracy (range of 81.1–99.5% across individual participants), speech and baseline with a mean of 57.0-78.9% (range of 32.9–92.5%). Participant-wise model accuracies over all states ranged from 58.9% to 91% (mean = 73.9%; ***Figure 3***B), all well above the 20% chance level for five classes. We also compared the classification performance between participants with right-hemisphere electrode coverage and those with left-hemisphere coverage. We found that the misclassification with baseline and speech was significantly more pronounced in the right-hemisphere group (see **Supplementary Figure 2** for accuracy and permutation tests), suggesting that differences in hemispheric language dominance could contribute to this specific classifier overlap. We then transformed the SVM weights to identify each state’s discriminative connections (***Figure 3***D) revealing that each state is represented by a small subset of connections (mean across states <≈ 0.002% of all possible connections) largely unique to that state (uniqueness ratio = 0.743; ***Figure 3***C, see Statistical Analyses). Spatially, auditory perception relied on STG–frontal links, visual states on occipital–frontal connections, and speech production on enhanced speechmotor cortex coupling as well as motor–auditory anti-correlations (***Figure 3***D and ***Figure 5***B). In contrast, a traditional analysis of strongest connections (i.e. correlation magnitude thresholding; see Single-Trial Functional Connectivity) yielded denser, overlapping connections (uniqueness = 0.283; ***Figure 3***C and E). These results show that single-trial FC decoding isolates a highly specific connectivity signature for each behavioral state, highlighting functionally meaningful interactions that are obscured when connectivity is addressed solely by correlation magnitude.

**Figure 3.**
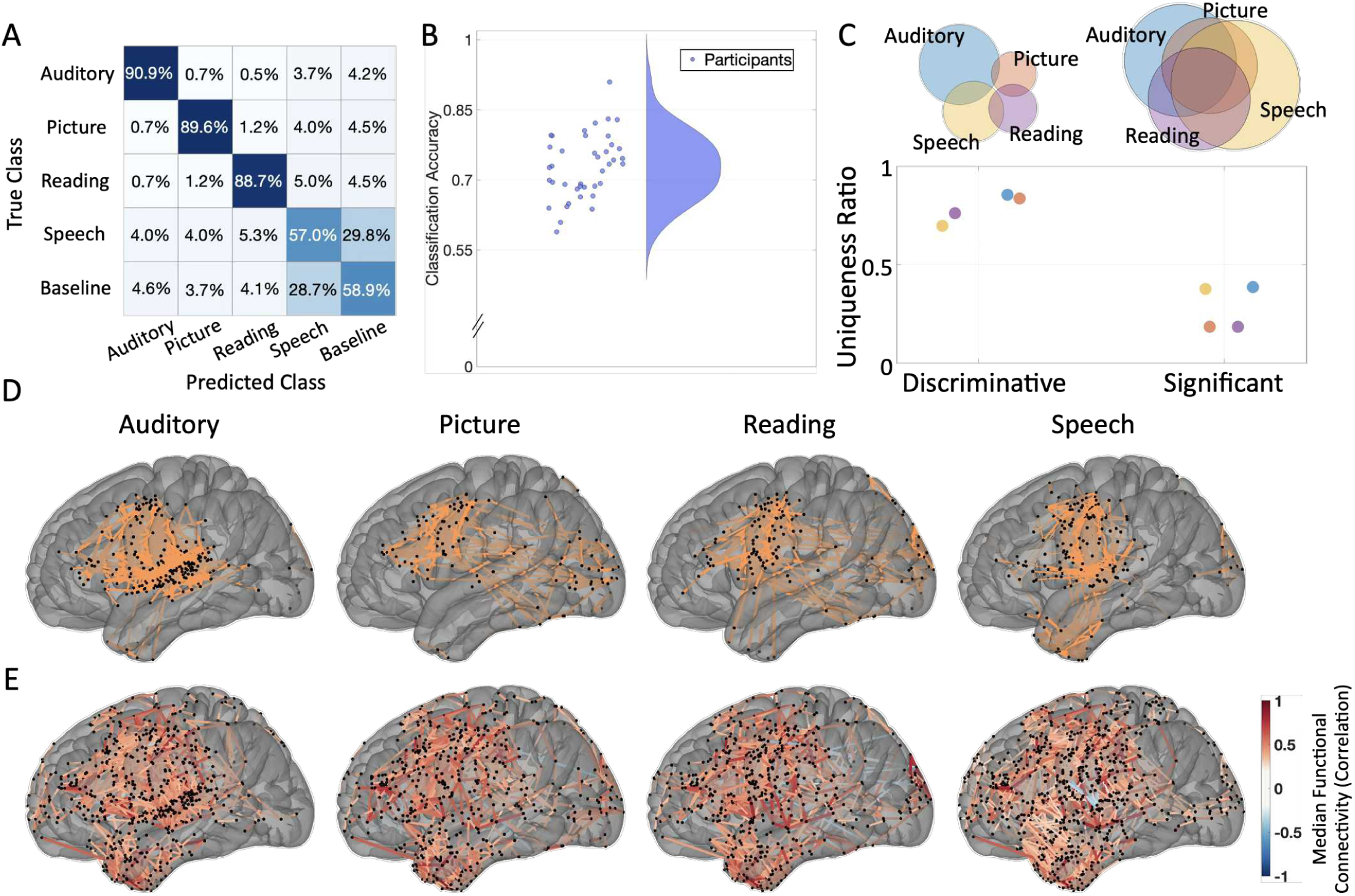
State-Specific Functional Connectivity patterns across all participants. (**A**) Confusion matrix obtained from five-fold cross-validated SVM classification, summed over 38 participants. Diagonal cells show per-class accuracies; off-diagonals show misclassification rates. (**B**) Distribution of participant-wise classification accuracies. The mean accuracy of 73.9% is well above the 20% chance level for five classes. (**C**) “Uniqueness ratio” for each behavioral state, comparing the sparsity of connections selected by our discriminative approach with those retained after a magnitude threshold (i.e. strongest connections). This ratio is closer to 1 when the connections are more unique in each state, and closer to 0 when they are overlaping across the behavioral states. Discriminative connections are significantly more state-specific than the strongest connections (average uniqueness ratio = 0.743, 0.283 respectively; two-sided permutation test, *p* < 0.0001 for all states). The venn diagrams on the top row show the distribution and overlap of the connections across states, with size of each circle representative of the number of connections across behavioral states. (**D**) Cortical distribution of discriminative connections for each state, across all participants. The patterns align with our current knowledge of involved regions in these states (e.g., STG-frontal links in auditory perception, motor–auditory decoupling in speech production). (**E**) Cortical distribution of connections selected solely by magnitude thresholding. The widespread, non-specific pattern contrasts with panel D, underscoring the greater specificity of the discriminative approach. (All brain surfaces are shown in left-hemisphere lateral view; separate right and left hemisphere results are shown in **Supplementary Figure 3**.)

### Relationship between discriminative connectivity and local neural activity

Speech production is notoriously difficult to study with non-invasive measures given the inherent motor artifacts. Conversely, intracranial studies have shown great promise in characterizing speech production dynamics, however an overwhelming number of studies have focused on local neural activity alone. We were interested in characterizing the unique contribution of our approach compared with the common practice of analyzing local high-gamma neural activity. During speech production, high-gamma active electrodes (see Statistical Analyses) on the cortex were predominantly distributed over the superior temporal and peri-central cortices (***Figure 4***A and E). This highly selective distribution has been a hallmark of local activity and is in stark contrast to the widespread distribution of electrodes across cortex that show a strong functional correlation (***Figure 4***B,D,E). Our decoding approach provides evidence for a more selective set of discriminative functional connections with broad coverage of the language network (***Figure 4***C,D,E). Our approach reveals a larger ratio of electrodes identified across inferior frontal and parietal cortices compared with local activity alone (permutation test, see Statistical Analyses). This indicates that local activity alone may be insufficient for identifying important regions in the language network and a connectivity-based approach is necessary as a complement.

**Figure 4.**
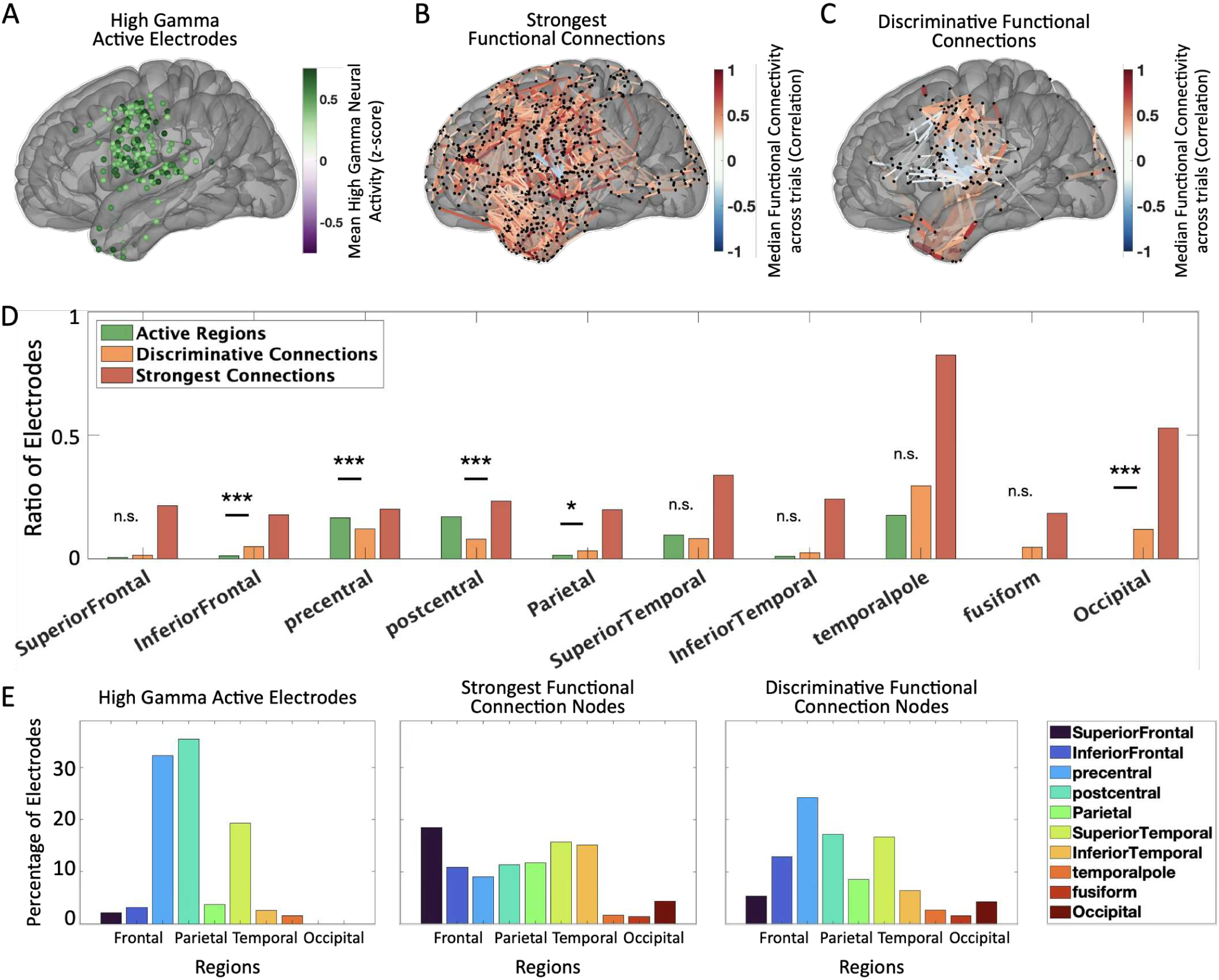
Comparison of maps for Speech production using three different methods. (**A**) Electrode-wise map of high-gamma activity amplitude (70–150 Hz) during speech production, aggregated across 38 participants and averaged across trials and time. (**B**) Connections whose median Pearson correlation exceeds a p *<* 0.05 threshold. This magnitude-based connectivity thresholding approach yields a dense, overlapping network. (**C**) Connections identified by the discriminative SVM transformed weights, resulting in a sparse and state-specific distribution of the connections. (**D**) Ratio of electrodes identified in each region compared to the total number of electrodes recording in that region, using three different approach. The lower ratios for ‘Active Regions’ and regions involved in ‘Discriminative Connections’ indicate that these methods are more sensitive in identifying relevant electrodes compared to the ‘Strongest Connections’ approach. Stars mark regions where the proportion for discriminative connections differs from active regions (permutation test, *p* values listed: Superior Frontal *p* = 0.27, Inferior Frontal *p* = 0.0004, Precentral *p* = 0.0004, Postcentral *p* = 0.0001, Parietal *p* = 0.0413, Superior Temporal *p* = 0.228, Inferior Temporal *p* = 0.167, Temporal Pole *p* = 0.0702, Fusiform *p* = 0.128, Occipital *p* = 0.0007). (**E**) Bar plots summarizing how identified electrodes are distributed among brain regions.

While our approach elucidates discriminative connections, we were interested in understanding the relationship between the degree of local activity and discriminative connectivity. To quantify this, we extracted the median correlation for all discriminative connections and the mean neural activity of their respective nodes (for every behavioral state; see Neural Activity versus Connectivity Analysis). The resulting connectivity values span both positive and negative correlations (***Figure 5***A), yet organize into clear, state-specific patterns when projected onto the cortex (***Figure 5***B). Node activity shows a similar broad distribution (***Figure 5***C) with a clear spatial gradient from low to high activity regions (***Figure 5***D). A natural question is whether nodes with higher neural activity contribute more to functional connectivity. To test this, we correlated the mean connectivity of each node in discriminative connections with its mean activity for every behavioral state and found significant correlations across states (Spearman Correlations, Auditory *r*ℎ*o* = 0.4472, *p* < 0.0001; Picture *r*ℎ*o* = 0.2925, *p* = 0.0003; Reading *r*ℎ*o* = 0.4306, *p* < 0.0001; Speech *r*ℎ*o* = 0.1195, *p* = 0.0254.). When visualizing the relationship (***Figure 5***E, color-coded by behavioral state), most nodes align along the diagonal, indicating balanced activity and connectivity. More active regions tend to exhibit stronger correlations. However, many electrodes diverge from this relationship: some are highly connected despite low neural activity, whereas others are strongly active but weakly connected. We quantified this imbalance with a connectivity–activity relative-distance index (RDI) and mapped it on the cortex (***Figure 5***F). The RDI highlights precentral and frontal gyri during auditory perception and speech-motor cortex during picture and reading perception, where connectivity exceeds activity. Conversely, during speech production discriminative nodes include both positive RDI (i.e. low activity and high connectivity) and negative RDI sites (i.e. high activity and low connectivity), particularly within the superior temporal, inferior frontal and precentral gyri. Together, these findings show that discriminative connectivity results in a unique, functionally meaningful network signature that is invisible to activity-only or correlation-magnitude-only views and underscores that connectivity adds a complementary dimension to local activity for differentiating behavioral states.

**Figure 5.**
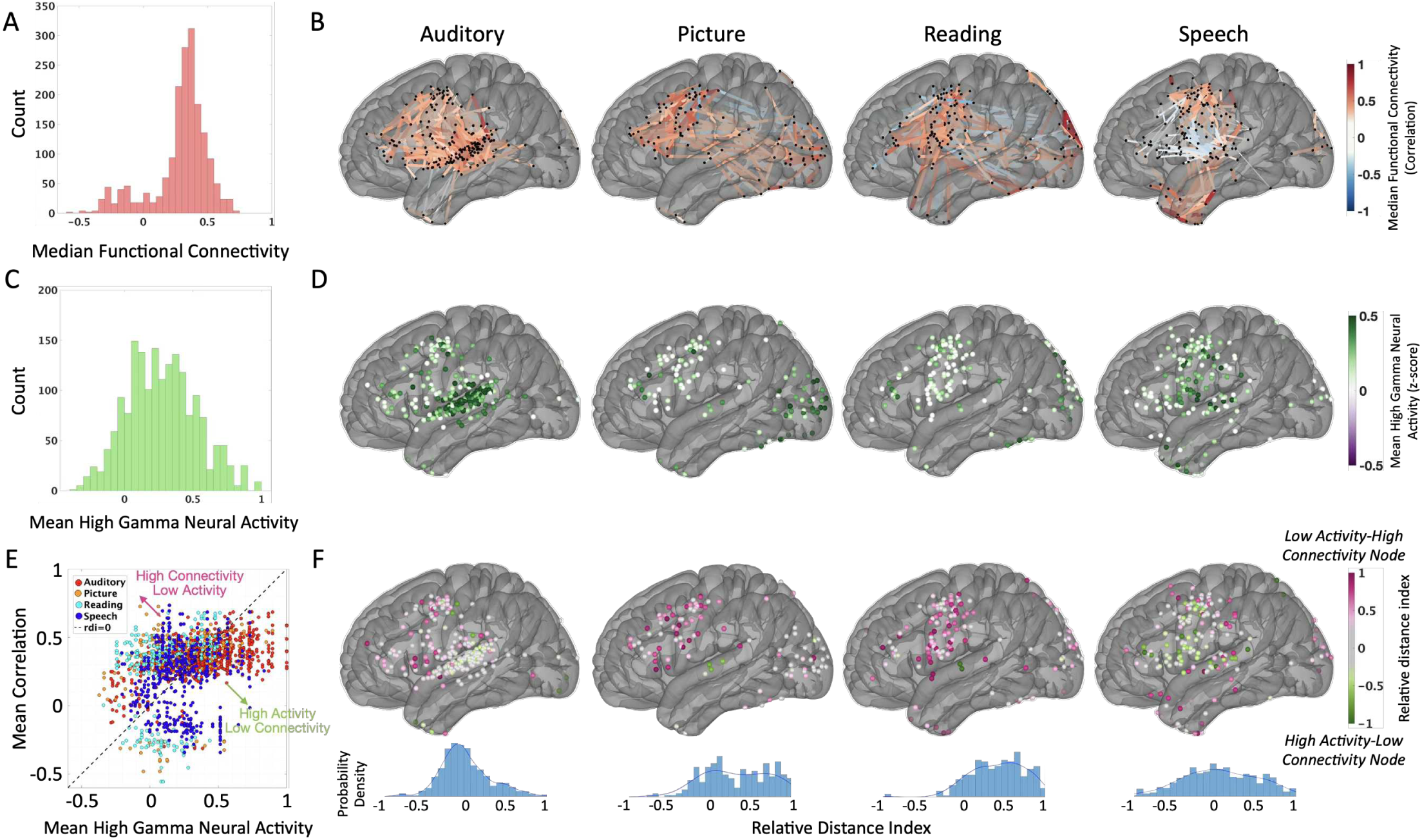
Relationship between discriminative connectivity and local high-gamma activity. (**A**) Histogram of FC values for all discriminative edges (pooled across states). While skewed toward stronger correlations, the distribution includes many weak connections that conventional magnitude thresholds would overlook. (**B**) The cortical distribution of state-specific discriminative connections. The color represents median functional connectivity values, representing the prominent connectivity patterns that remain stable across the trials of these state. (**C**) Histogram of high-gamma activity, averaged across trials and time at nodes participating in discriminative connections. (**D**) The cortical distribution of mean high-gamma activity, only for nodes participating in discriminative connections of each state. (**E**) Scatter-plot of mean connectivity strength versus mean high-gamma activity for the nodes participating in discriminative connections. A positive trend is present, yet many nodes deviate markedly, indicating regions with high connectivity but low signal activity and vice versa. (**F**) Connectivity–activity relative distance index projected onto the cortex (pink = nodes with relatively higher connectivity than activity; green = nodes with higher activity than connectivity). This map highlights regions whose functional importance is underestimated by activity-only analyses. (All brain surfaces are shown in left-hemisphere lateral view; separate right and left hemisphere results are shown in **Supplementary Figure 4**.)

## Discussion

Our study leverages single trial connectivity decoding across a large cohort of patients (N = 38) to investigate functional connectivity. Our approach is able to achieve high decoding across five speech-related behavioral states while providing interpretable weights that highlight the discriminative connections. Compared to conventional methods (i.e. local high-gamma activity), our approach reveals previously obscured cortical sites that discriminate behavioral states. These results are among a select few functional connectivity studies focusing on speech related cortical networks, providing a rare window into the functional networks supporting this function.

Despite valuable insights, most human FC studies in ECoG share design choices that blur fast connectivity dynamics. Many studies estimate connectivity from long windows of >1s (***Kucyi et al., 2018***; ***Nir et al., 2008***). Investigating connectivity during the transient behavioral states involved in speech perception and production will require a shorter time window, enabling us to capture these state specific characteristics of connectivity. Further, studies often average the signal or connectivity over many repetitions, which eliminates trial-to-trial variability (***Casimo et al., 2019***; ***Foster et al., 2015***). Averaging signals or correlation matrices can result in robust estimations that highlights the strongest connections. However, this step also blurs the distinction between state specific signals or connections that maintain a steady low value and those that vary drastically. To be able to capture the transient, state-specific connections that are stable across trials regardless of their strength, we compute FC in 500-ms windows and keep every trial separate. This allows us to preserve within-state variability while avoiding the loss of temporal detail. As a result, this framework unveils the fast dynamics of speech perception and production with a resolution that respects both the transient and distributed nature of these networks.

To investigate the discriminative connections underlying speech perception and production, we computed functional connectivity between every pair of electrodes. By leveraging the interpretability of the transformed weights of a linear SVM model, we detect sparse state-specific connections based solely on their ability to differentiate behavioral states (***Figure 3***D, ***Figure 5***B). As a result, we identify both strong and weak discriminative connections, including ones between regions that fall outside the canonical language areas. In contrast, the standard connectivity magnitude thresholding method results in a widespread and state-agnostic set of connections, masking these distinctions (***Figure 3***E). A common practice in ECoG-based FC studies that avoids this problem is to restrict analyses to a handful of predefined nodes using seed-based or region-of-interest strategies (***Keller et al., 2013***). Most often, these seeds are selected based on electrodes with large high-gamma responses or those deemed critical by electrical stimulation mapping (***Kingyon et al., 2015***; ***Rolston and Chang, 2018***; ***Fox et al., 2020***; ***Hsieh et al., 2024***). In the context of language, seed-based strategies typically target inferior-frontal and auditory cortices, privileging well-established nodes but marginalising others, like middle frontal, precentral, postcentral and temporal pole. While methodologically convenient, this choice significantly narrows the scope of network inference and implicitly reinforces the dominant cortical regions. Moreover, it overlooks potentially informative and functionally relevant regions with lower signal (activity or connectivity), such as those identified by our RDI close to 1 or −1 in ***Figure 5***F. Our results suggest that a more inclusive (e.g. all electrodes) approach coupled with a decoding framework reveals a set of meaningful connections and avoids anatomical blind spots.

Our findings reveal connections across peri-sylvian cortices as well as the temporal pole during shifts from perception to production — a dynamic that has been difficult to capture with previous approaches. While fMRI studies have been instrumental in mapping large-scale networks (***Biswal et al., 1995***; ***Smith et al., 2009***; ***Fox et al., 2005***; ***Greicius et al., 2003***; ***Power et al., 2011***; ***Yeo et al., 2011***) and in characterizing the language network (***Saur et al., 2008***; ***Braga et al., 2020***; ***Tomasi and Volkow, 2012***; ***Branco et al., 2020***; ***Lipkin et al., 2022***), most task-based studies focus on listening, reading, or covert speech. This has left overt speech underexplored in fMRI, due to motion artifacts (***Birn et al., 1999***; ***Gracco et al., 2005***). Moreover, the temporal smoothing of the BOLD response limits fMRI’s ability to resolve the rapid state transitions between perception and production of speech. In contrast, ECoG, with its millisecond resolution, allows us to track these fast dynamics. Our approach captures connectivity patterns directly relevant to speech processes, revealing perception-to-production shifts with millisecond resolution. Specifically, we find positive correlations during perception across frontal and relevant sensory regions, however speech production is associated with peri-rolandic and temporal (superior temporal gyrus and anterior temporal lobe) networks. Interestingly, the discriminative connections for speech production included many negative correlations (***Figure 5***B) between superior temporal gyrus and ventral frontal cortices (inferior frontal gyrus and ventral pre-central gyrus) which likely reflects speech induced suppression during vocalization (***Khalilian-Gourtani et al., 2024***; ***Kurteff et al., 2024***). Although ECoG coverage is clinically constrained, it complements the whole-brain approach of fMRI by offering a high-resolution view of speech processing dynamics at behaviorally relevant timescales. The present cohort consisted exclusively of subdural grid and strip electrodes, which limits our view to neocortical surface regions. Future work incorporating depth or stereo-EEG (sEEG) electrodes would enable examination of subcortical and mesial temporal contributions (including hippocampus and amygdala) to speech-related connectivity networks, which remain inaccessible with surface recordings alone.

An important finding in our current results is the robust discriminative connectivity between anterior temporal lobe and peri-sylvian sites (e.g. dorsal precentral and inferior parietal). This connectivity (***Figure 4***, ***Figure 5***, **Supplementary Figure 5E**) was predominantly seen during the speech production state and likely reflects word retrieval (***Ralph et al., 2017***). These speech related connections were mostly limited to zero lag connectivity (Supplementary Figure 6). We do not believe that these anterior temporal sites have any residual speech-related articulatory artifacts (see **Supplementary Figure 5** for a comparison and frequency profile of artifacts, and **Supplementary Figure 7** for a representative single-participant example contrasting single-trial activity and connectivity at included versus excluded ATL electrodes). These finding are consistent with reports of ATL activity during speech production (***Kurteff et al., 2024***; ***Ozker et al., 2022***; ***Morgan et al., 2026***; ***Abel et al., 2016***), however previous studies did not focus on this activity, likely due to limited coverage and number of sampled patients. We believe that the prominence of this ATL pattern during speech likely supports lexical access as part of a multi-modal semantic hub (***Ralph et al., 2017***).

Our results further show that discriminative connections emerge in regions with both high and low neural activity. In the ECoG literature, neural activity is most often measured as the amplitude of the high-gamma (70–150 Hz) frequency range, with task-relevant regions mapped by significant increases above baseline. Numerous studies have shown that high-gamma active regions during speech tasks localize strongly to IFG, precentral gyrus, STG, and MTG (***Chang et al., 2010***; ***Edwards et al., 2010***; ***Flinker et al., 2015***; ***Moses et al., 2016***). This approach maps active electrodes but often overlooks distributed connectivity as well regions that are not significantly active. Our connectivity findings provide evidence for connections involving low-activity electrodes, particularly in parts of precentral and inferior frontal cortices during perceptual states (***Figure 5***F). The fact that we find these connections in addition to discriminative connections between electrodes with high neural activity, suggests that the functional networks supporting speech extend beyond the most metabolically active regions. The distribution of these sites predominantly span prefrontal cortices (see **Supplementary Figure 8**) and are in line with recent findings providing evidence for linguistic information representations encoded with low neural activity (***Morgan et al., 2026***). It is likely that these high connectivity values reflect interconnected cortical regions that represent linguistic and state-specific information despite low neural activity. Further, the identification of both active-to-active and active-to-silent discriminative connections demonstrates that connectivity analyses provide orthogonal information to local high-gamma measures, potentially revealing preparatory or coordinating regions that studies relying on activation alone may overlook. This orthogonal dimension might explain why some sites identified as causally necessary for function (via clinical stimulation mapping) do not necessarily exhibit increased local activity. Connectivity may be a more reliable measure for predicting clinically relevant language sites identified by electrical stimulation mapping (***Emami et al., 2026***; ***Hsieh et al., 2024***).

Together, these findings highlight how single-trial functional connectivity decoding provides a powerful complement to traditional analyses of neural activity. By uncovering state-specific and sparse connections across regions, we reveal a distributed speech network that cannot be captured by focusing on local activation alone or seed-based connectivity. The ability to decode behavioral states from transient connectivity patterns underscores the dynamic and flexible nature of the underlying cognitive processes and allows for mapping the full cortical networks involved in speech perception and production.

## Methods and Materials

### Participant Information

We used intracranial data from 38 neurosurgical patients undergoing surgical monitoring for Epilepsy (17 females, mean age: 30.82 ± 12.2 yo, 15 right grid, 23 left grid coverage). Patients were selected based on coverage of electrodes sampling from the patients’ coverage across the superior temporal, precentral, postcentral, and inferior temporal gyri, and their ability to perform the experimental tasks. No minimum electrode count threshold was applied; inclusion was based on qualitative assessment of spatial coverage and signal quality (A summary of brain regions sampled is included in **Supplementary Table 2**). The electrode arrays consisted exclusively of subdural grids and strips placed directly on the cortical surface; no depth or stereo-EEG (sEEG) electrodes were included in the analysis. The implantation and location of the electrodes were guided solely by clinical requirements. All participants provided informed consent following approval from the NYU Langone institutional review board.

### Experimental Tasks and Behavioral States

Participants completed three tasks that elicited production of the same target words from auditory or visual stimuli (***Figure 1***A). The tasks were auditory word repetition (repeat an auditorily presented word), visual word reading (read a visually presented written word), and picture naming (name a word from a visually presented line drawing). Each task comprised 100 trials drawn from an identical set of 50 words that appeared twice across different modalities (The list of stimuli and associated number phonemes and syllables are included in **Supplementary Table 1**). For each task and trial, the stimulus was randomly presented. Each trial began with a silent baseline of at least 500 ms, followed by stimulus presentation, and then speech production (***Figure 1***C). Participants are instructed to produce the word as soon as they are ready without a go signal (***Shum et al., 2020***; ***Yu et al., 2025***). When participants finished articulation, the experimenter pressed a keyboard button to advance to the next trial, after-which a software random delay and an additional 500 ms delay were introduced to ensure a silent baseline period (Supplementary Figure 9). We defined five behavioral states using 500 ms windows time-locked to event markers: auditory perception (auditory stimulus onset), picture perception (picture stimulus onset), reading perception (written word stimulus onset), speech production (speech onset; shared across tasks), and baseline (pre-stimulus presentation period, also shared across tasks).

### Electrode Localization

Electrode localization in both individual and MNI space was performed by co-registering a preoperative structural MRI with a postoperative structural MRI or CT scan using a rigid-body transformation. To correct for post-implantation, edema-induced brain shifts, electrode coordinates were projected to the preoperative segmented cortical surface following established procedures (***Yang et al., 2012***). Non-linear registration to MNI space was achieved using the DARTEL algorithm (***Ashburner, 2007***). Individual anatomical labeling and assignment of electrodes to specific regions of interest (ROIs) were determined via automated FreeSurfer segmentation of each participant’s preoperative MRI. In this study, we focused exclusively on subdural grid and strip contacts across the N = 38 participants, spanning the lateral temporal, frontal, parietal, and lateral occipital cortices. All the plots and visualizations on the standard brain were implemented using the Mithra toolbox (***Khalilian-Gourtani et al., 2025***).

### ECoG Data Acquisition and Preprocessing

ECoG neural signals were recorded at 2048 Hz with the Neuroworks Quantum Amplifier (Natus Biomedical, Appleton, WI) and were downsampled to 512 Hz during export. Electrodes exhibiting seizure activity or excessive noise identified clinically were excluded from further analysis. Data were re-referenced using a common average referencing scheme to mitigate common-mode noise. We band-passed signals into eight logarithmically spaced sub-bands from 70–150 Hz and averaged them to isolate high-gamma activity. We adopted this multi-band extraction to avoid dominance of lower frequencies (***Hamilton et al., 2021***; ***Moses et al., 2016***). We computed analytic amplitude as the absolute value of the Hilbert transform of the filtered signal in each band. We then normalized the data (i.e., z-score) relative to the entire recording session, and subsequently averaged the normalized signals across bands. As an extra precautionary step against momentary artifacts, in each electrode, we identified trials whose high-gamma envelope had an out-of-distribution maximum or variance across trials. A trial was removed from the analysis if at least one electrode of the pair included an out of the distribution trial.

### Single-Trial Functional Connectivity

For computing functional connectivity, we segmented the ECoG high-gamma signals into 500 ms epochs for each trial and behavioral state, aligned to task event markers described above. For every epoch, we computed pairwise Pearson correlations of high-gamma analytic amplitude between all electrodes, producing a symmetric functional connectivity matrix per trial and state. To avoid redundancy, we vectorized the upper-triangular off-diagonal of each matrix to obtain a trial-specific connectivity vector for each state. Each vector contains the correlation for all possible electrode pairs (features), with the sign preserved. In the analysis for **Supplementary Figure 6** including different lags, we computed the pairwise Pearson correlation between the zero lag signal and the lagged (or leading) signal, creating a non-symmetrical connectivity matrix. We vectorized the matrices removing the diagonal (self connections), resulting in a feature space double the size of the original analysis.

In the analysis to identify the strongest connections (correlation magnitude-thresholding, ***Figure 3***E), we first summarized each state’s connectivity matrix by computing the median value of each connection across trials. We then evaluated how much each connection’s median deviated from the overall distribution of medians. Connections that showed significantly high or low values (beyond what we would expect by chance, after accounting for multiple comparisons, *α* < 0.05) were retained. All others were set to zero. After this thresholding step, we reconstructed the participant- and state-specific connectivity matrices using only the significant connections. Finally, we combined these matrices into a block-diagonal group matrix, preserving individual participant structure while enabling pooled network analysis.

### Decoding Framework

We classified trial-specific connectivity vectors into behavioral states using a linear Support Vector Machine (SVM) in MATLAB (MathWorks). To address class imbalance, we applied the Synthetic Minority Oversampling Technique (SMOTE, ***Chawla et al. (2002)***) within each participant and training fold to avoid leakage. We evaluated performance with stratified five-fold cross-validation per participant, averaged accuracy across folds, and derived confusion matrices. We estimated chance levels as theoretical chance (20% for five states, after accounting for SMOTE) and verified this estimate with a permutation approach (i.e. shuffling state labels 10000× within participant). To identify discriminative connections in ***Figure 2*** and ***Figure 3***D, we transformed SVM weights from the decision boundary hyperplanes using the Haufe forward model (***Haufe et al., 2014***). This transformation enables interpretation of classifier weights by accounting for feature dependencies and mapping them back to neural sources. We normalized the resulting weights using their *L*2 norm, and then evaluated how far each transformed weight deviated from the median of the distribution, using the median absolute deviation (MAD) as a robust scale estimator. We set to zero all weights whose deviation from the median did not exceed a threshold derived from a Laplacian tail bound (*α* < 0.05). We selected this threshold to reflect the heavy-tailed distribution of weights. We defined discriminative connections as those whose weights exceeded this threshold, indicating they contributed most strongly to the separation of behavioral states in the classifier. To quantify the specificity of the identified connections for each state, we computed a uniqueness ratio. This ratio represents the proportion of connections uniquely associated with a given behavioral state relative to all connections identified for that state.

### Neural Activity versus Connectivity Analysis

For the analysis in ***Figure 5***, we focused on electrodes that participated in the discriminative connections. For each such electrode and state, we quantified local activity as the mean high-gamma analytical amplitude across trials and time points. For each discriminative connection, we quantified connectivity strength as the median Pearson’s correlation across trials of that state. We then computed a connectivity–activity relative distance index (RDI) at the electrode level:

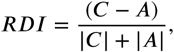

Where A denotes average local activity and C denotes average connectivity. RDI ranges from −1 (high activity, low connectivity) to +1 (low activity, high connectivity), with values near zero indicating balance. We projected RDI onto each participant’s cortical surface to visualize its spatial distribution.

### Statistical Analyses

For assessing the significant weights and connections (i.e. Decoding Framework and Single-Trial Functional Connectivity, respectively), We performed two-sided hypothesis tests at the single participant level. To control the family-wise error rate (FWER) across multiple comparisons, we applied a Bonferroni correction, maintaining an FWER of *α* = 0.05. For active electrode selection, we compared the high-gamma neural activity of each trial averaged over time, with the distribution of high-gamma neural activity during baseline, and used a one-sided threshold with *α* = 0.05 of the baseline to label an electrode as active on that trial.

To compute the significance of the difference between uniqueness ratios (***Figure 3***C), we performed permutation test, randomly swapping the connections identified across the two methods (for each connection, values are 1 for significant/discriminative connection and 0 otherwise, then swapped between methods with probability 0.5), with 10,000 permutations. We controlled for multiple comparisons using the false-discovery rate for a *p* < 0.05 two-sided significance (***Benjamini and Hochberg, 1995***).

To identify brain regions where the two methods yielded different results (***Figure 4***D), we compared the fraction of electrodes identified by the discriminative FC method against the fraction identified by the neural signal-only method. For each state *s* and region *r*, we defined the fraction(ratio) as the number of identified electrodes in *r* divided by the total electrodes recorded in *r*. This was statistically compared to a null hypothesis distribution by randomly permuting region labels across electrodes (preserving each region’s total electrode count) while holding fixed which electrodes were identified by that method, recomputing the ratio difference 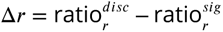 in each region for 10,000 permutations. Two-sided p-values used the permutation tail; For each region and state, we controlled for multiple comparisons using the false-discovery rate (***Benjamini and Hochberg*** (***1995***), *α* = 0.05).

## Acknowledgments

This work was supported by National Institutes of Health grants R01NS109367, R01NS115929, and R01DC018805 (A.F.) and National Science Foundationshum IIS-2309057.

## Supplementary Information

**Supplementary Figure 1.**
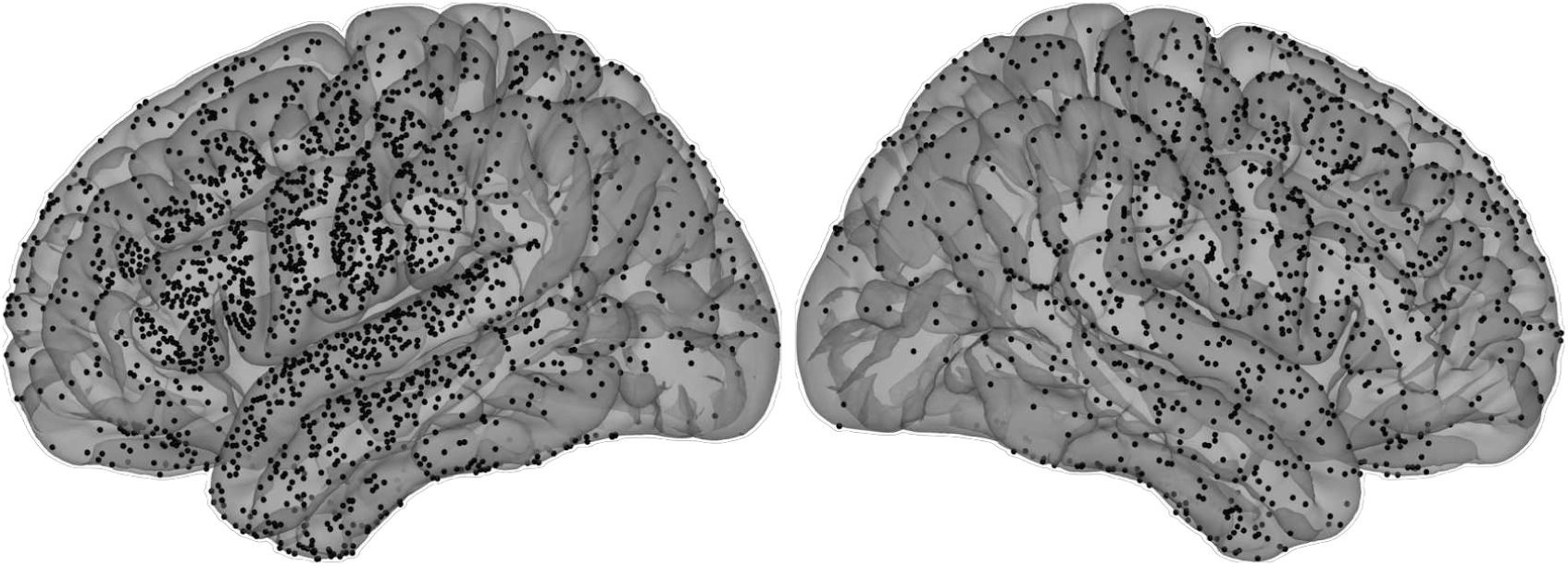
The spatial electrode coverage across participants, showing both right and left hemisphere.

**Supplementary Figure 2.**
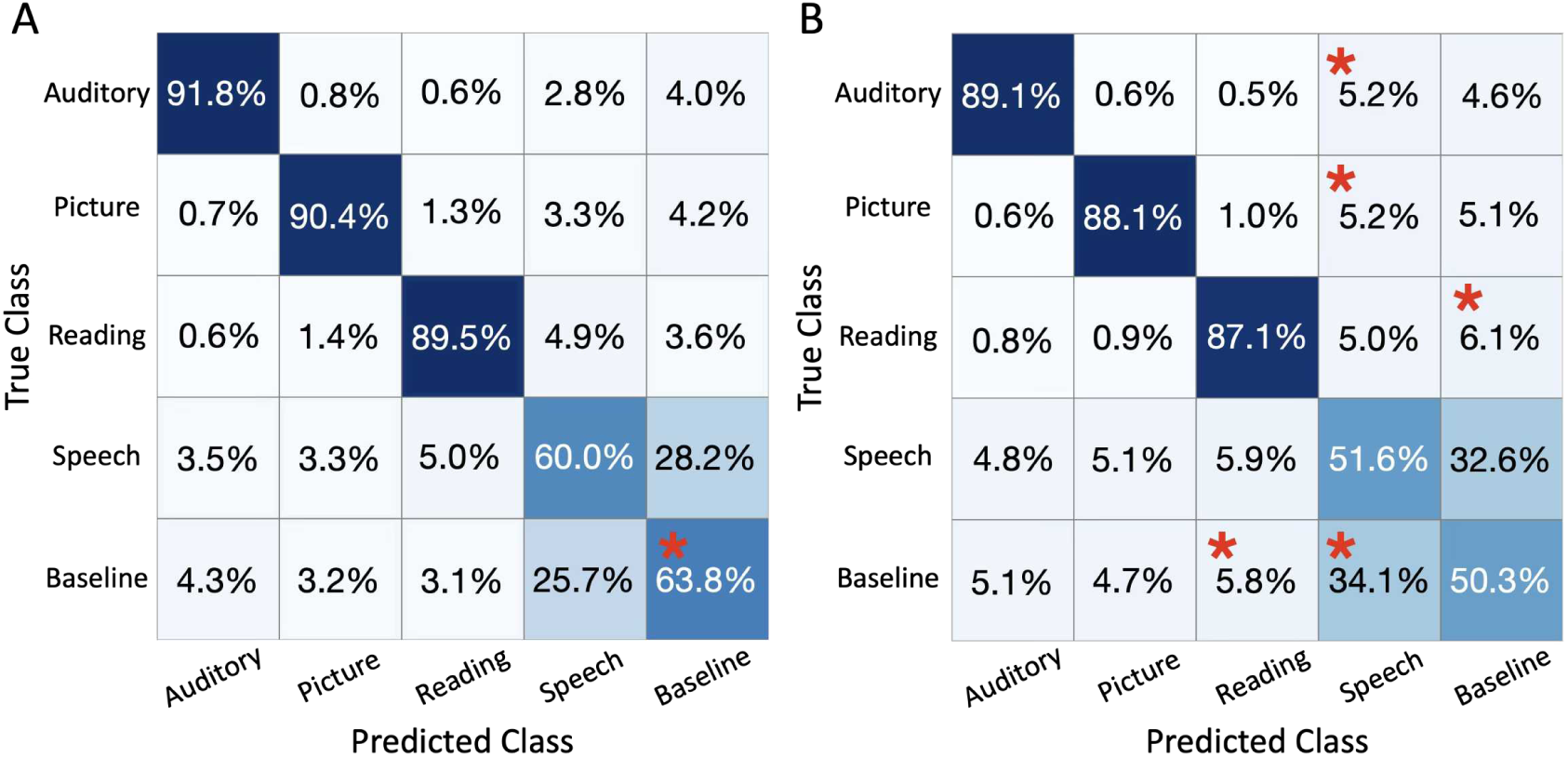
Behavioral state classification confusion matrices by hemisphere. Confusion matrices showing five-fold cross-validated SVM classification accuracy for participants with electrodes implanted in the left hemisphere (panel A) and right hemisphere (panel B). Rows indicate the true behavioral state, and columns indicate the predicted class. Red asterisks (*) denote cells where classification rates differ significantly between the left and right hemisphere groups (permutation test, N = 10000, p < 0.05). Right hemisphere coverage shows increased misclassification of perception states as speech or baseline, as well as higher mislabeling of baseline trials as speech.

**Supplementary Figure 3.**
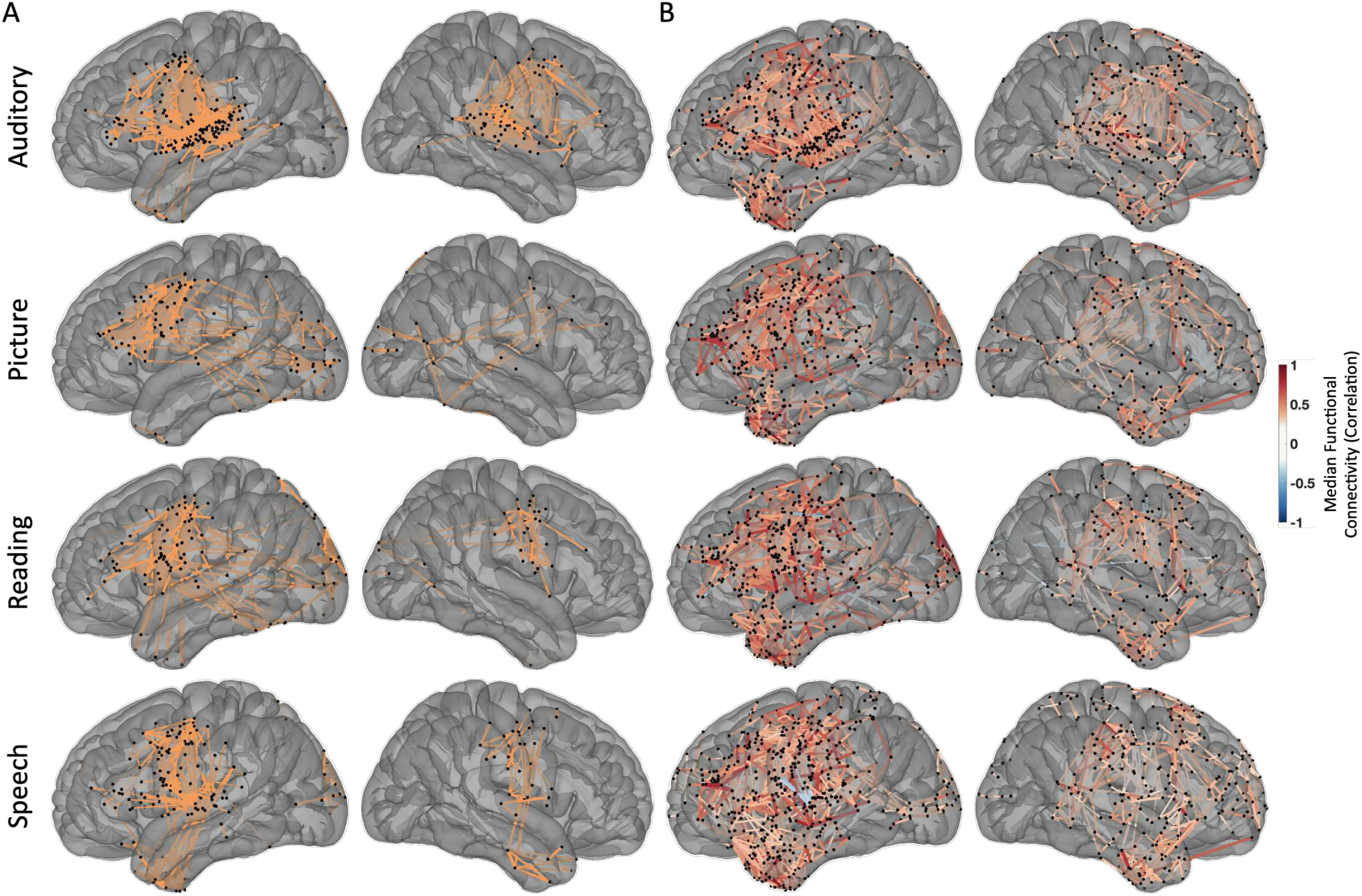
Functional Connections identified using two methods, separated for participants with *left hemisphere* and *right hemisphere* coverage. (A) Cortical distribution of discriminative connections for each state. Orange lines depict connections whose median absolute deviation of the transformed SVM weights survive a Laplacian tail bound significance threshold; node positions correspond to electrodes. (B) Cortical distribution of connections selected solely by magnitude thresholding. The widespread, non-specific pattern contrasts with panel A, underscoring the greater specificity of the discriminative approach.

**Supplementary Table 1.** Stimuli information and associated number phonemes and syllables. Index is matched with picture naming dataset index.

| Index | Word | Number of Phonemes | Number of Syllables |
| --- | --- | --- | --- |
| 2 | airplane | 6 | 2 |
| 8 | arrow | 3 | 2 |
| 14 | ball | 3 | 1 |
| 15 | balloon | 5 | 2 |
| 22 | bed | 3 | 1 |
| 23 | bee | 2 | 1 |
| 25 | bell | 3 | 1 |
| 35 | box | 4 | 1 |
| 36 | bread | 4 | 1 |
| 39 | bus | 3 | 1 |
| 41 | button | 5 | 2 |
| 42 | cake | 3 | 1 |
| 49 | cat | 3 | 1 |
| 67 | couch | 3 | 1 |
| 68 | cow | 2 | 1 |
| 69 | crown | 4 | 1 |
| 73 | dog | 3 | 1 |
| 80 | drum | 4 | 1 |
| 81 | duck | 3 | 1 |
| 85 | envelope | 7 | 3 |
| 90 | flag | 4 | 1 |
| 97 | fork | 4 | 1 |
| 105 | glasses | 6 | 2 |
| 106 | gloves | 5 | 1 |
| 109 | grapes | 5 | 1 |
| 112 | gun | 3 | 1 |
| 114 | hammer | 4 | 2 |
| 118 | hat | 3 | 1 |
| 119 | heart | 4 | 1 |
| 122 | house | 3 | 1 |
| 128 | key | 2 | 1 |
| 129 | kite | 3 | 1 |
| 130 | knife | 3 | 1 |
| 131 | ladder | 4 | 2 |
| 134 | leg | 3 | 1 |
| 143 | lock | 3 | 1 |
| 146 | moon | 3 | 1 |
| 149 | mouse | 3 | 1 |
| 155 | nose | 3 | 1 |
| 168 | pencil | 6 | 2 |
| 202 | sheep | 6 | 2 |
| 211 | socks | 4 | 1 |
| 226 | table | 5 | 2 |
| 235 | toe | 2 | 1 |
| 245 | umbrella | 7 | 3 |
| 246 | vase | 3 | 1 |
| 248 | violin | 6 | 3 |
| 250 | watch | 3 | 1 |
| 253 | well | 3 | 1 |
| 257 | window | 5 | 2 |

**Supplementary Figure 4.**
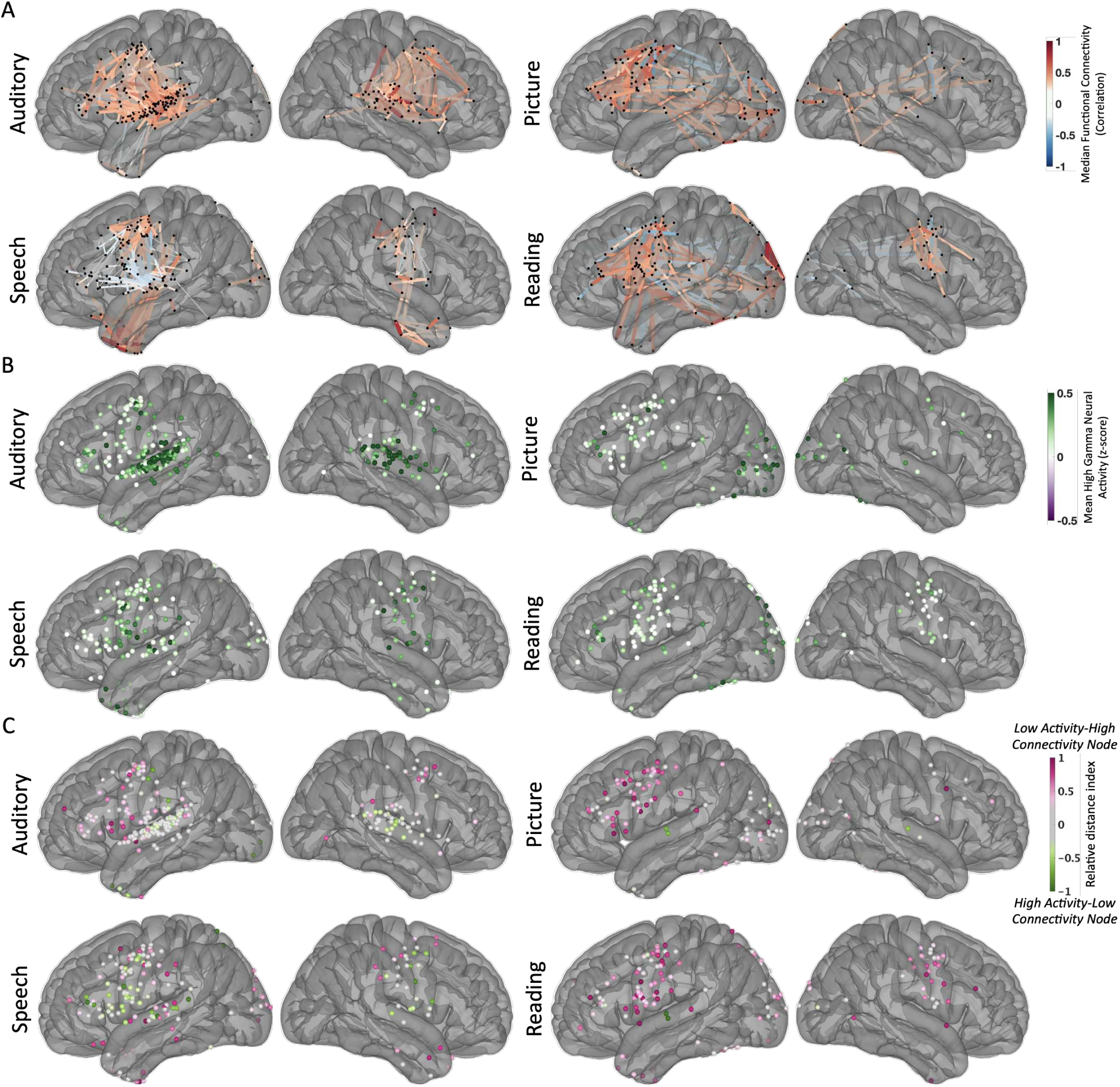
Relationship between discriminative connectivity and local high-gamma activity, separated for participants with *left hemisphere* and *right hemisphere* coverage. (A) The cortical distribution of state-specific discriminative connections, visualized on the right lateral view of an average brain. The color represents median functional connectivity values, representing the prominent connectivity patterns that remain stable across the trials of these state. (B) The cortical distribution of mean high-gamma activity, only for nodes participating in discriminative connections of each state. (C) Connectivity–activity relative distance index projected onto the cortex (pink = nodes with relatively higher connectivity than activity; green = nodes with higher activity than connectivity). This map highlights regions whose functional importance is underestimated by activity-only analyses.

**Supplementary Figure 5.**
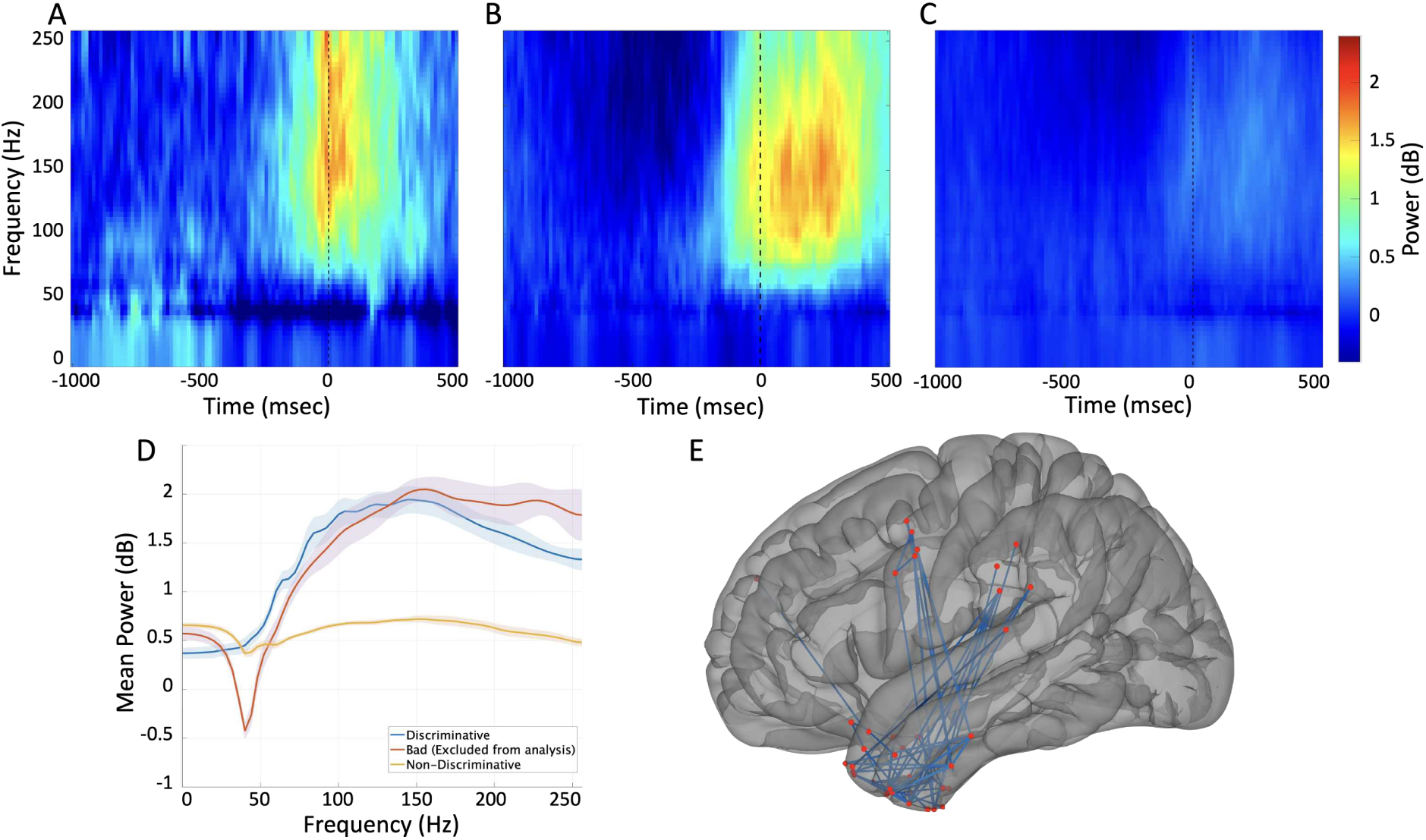
Time-frequency dynamics of ATL electrodes, grouped by inclusion status and discriminability for speech. (A) Mean spectrogram (time-frequency power, baseline-normalized in dB) for ATL electrodes that were excluded from analysis due to motion artifacts. (B) Mean spectrogram for ATL electrodes that were included in the analysis and showed at least one discriminative functional connection for the speech class. (C) Mean spectrogram for all remaining included ATL electrodes that did not show discriminative connectivity for speech (non-discriminative). In A-C, the dashed vertical line at 0 ms marks speech onset, and color indicates power change (dB) relative to baseline. (D) Mean power spectra (power averaged over electrodes and a time window from −50 to +100 ms relative to speech onset) as a function of frequency, shown for each of the three electrode groups: bad/excluded (red), discriminative (blue), and non-discriminative (yellow), corresponding to the electrode sets shown in A, B, and C, respectively. The shaded colors exhibit standard error across electrodes on each group.

**Supplementary Figure 6.**
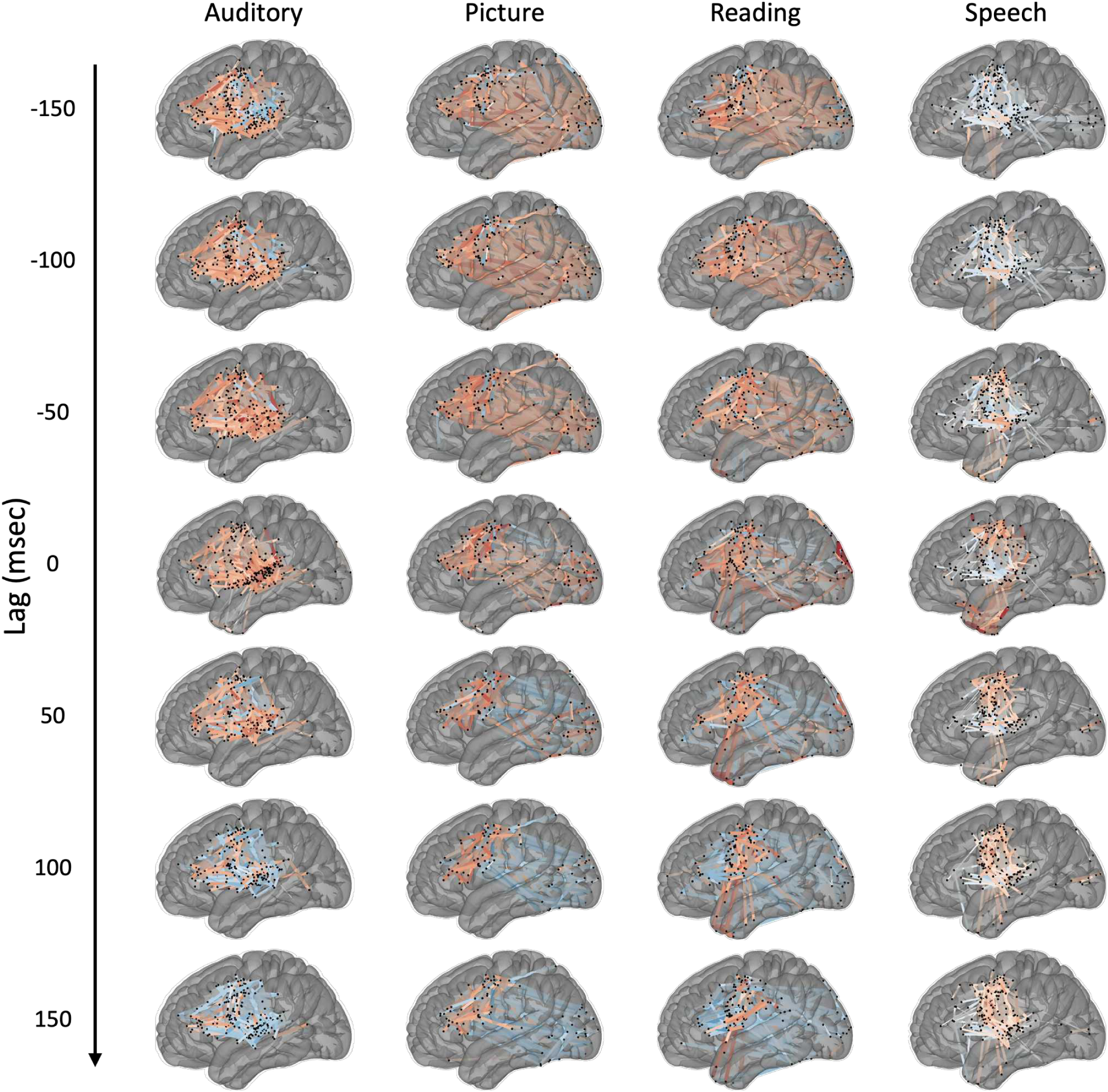
Robustness of discriminative connections to epoch time lag (or lead). Median functional connectivity across trials for each of the four cognitive states (Auditory, Picture, Reading, and Speech, represented in columns). The results are computed after shifting the analysis epoch by a fixed lag relative to original timestamps, from −150 ms (epoch shifted earlier) to +150 ms (epoch shifted later) in 50 ms steps (rows). For each lag, the epoch window used to extract each state’s data was shifted uniformly by that amount, and pairwise electrode correlations, feature extraction, and classification were repeated as in the main analysis. Black dots indicate electrode locations; lines indicate connectivity weights surviving thresholding, colored by sign (warm colors, positive; cool colors, negative) and scaled by magnitude. Connectivity topographies remain qualitatively similar across the full shift range for each state, indicating that the reported patterns are not an artifact of the specific epoch boundaries chosen and are stable over this temporal window.

**Supplementary Figure 7.**
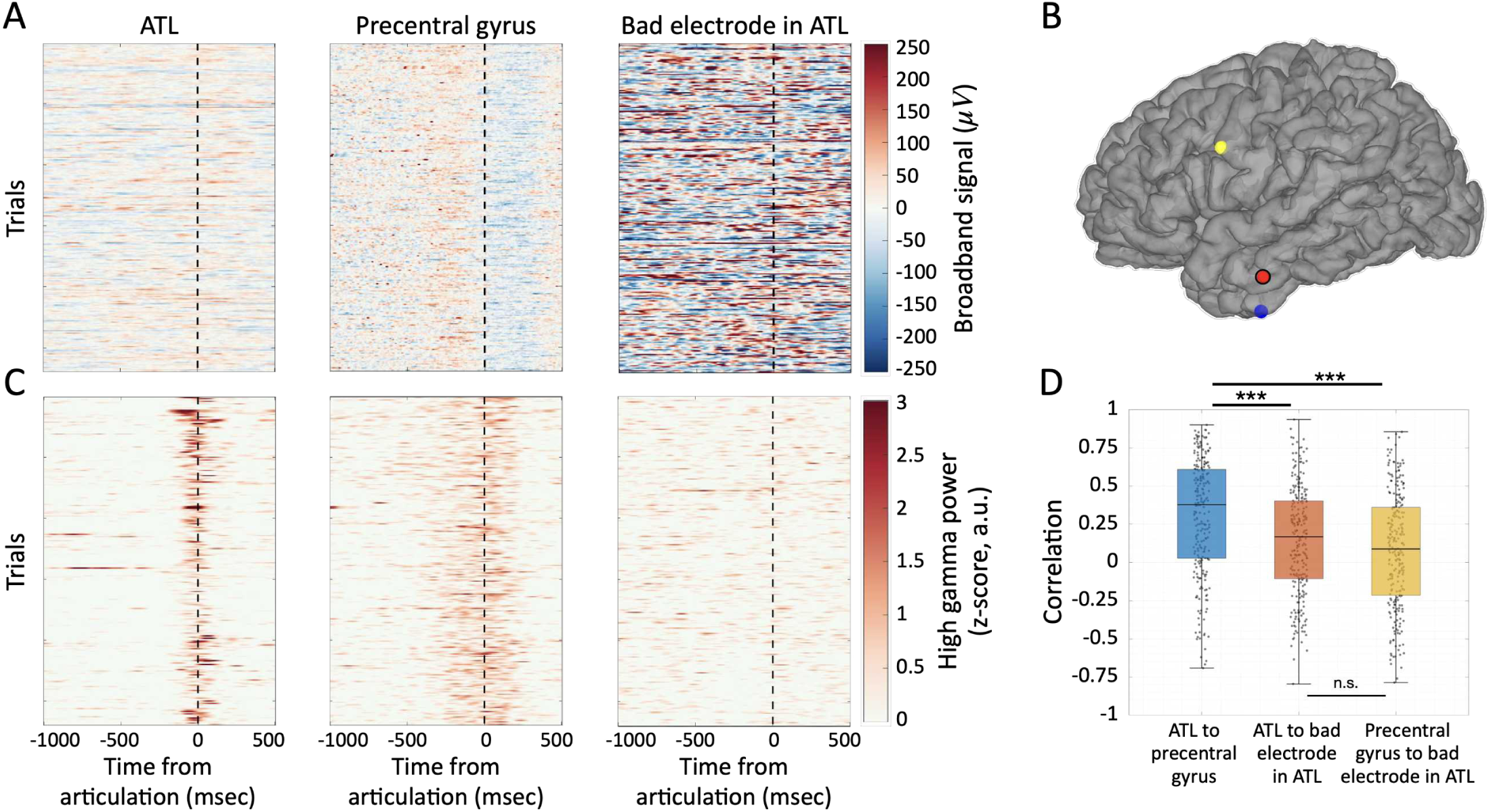
Single-trial activity and connectivity for an included ATL electrode versus an excluded ATL electrode, in a representative participant. All panels are computed from the same set of N = 252 speech-production trials, response-locked to articulation onset (dashed line).(A) Single-trial broadband (unfiltered, common-average-referenced) traces for three electrodes: an anterior temporal lobe (ATL) electrode forming a discriminative speech connection (blue in B), the precentral gyrus electrode to which it is connected in that discriminative connection (yellow in B), and an ATL electrode excluded from analysis by our EMG/artifact criteria (red in B, with black outline). each row represents a trial and are in chronological order, with the same ordering applied to all columns in A and C. All three panels share the color scale. (B) Electrode locations on the participant’s reconstructed cortical surface; the excluded electrode (red) is marked with a black outline. (C) Single-trial high-gamma power (70–150 Hz, z-scored) for the same three electrodes and trial ordering as in A. The two included electrodes (ATL, precentral) show consistent articulation-locked high-gamma responses; the excluded electrode shows no event-related high-gamma activity. (D) Distributions of trial-wise high-gamma functional connectivity (Pearson correlation) for the three electrode pairs; each dot is one trial, boxes show median and interquartile range. The discriminative ATL–precentral connection (blue) is significantly more correlated than either connection to the excluded electrode (ATL–excluded, red; precentral–excluded, yellow). *** p < 0.001, paired permutation test (10,000 permutations), Bonferroni-corrected across the three comparisons; n.s., not significant. The two connections involving the excluded electrode do not differ from each other.

**Supplementary Figure 8.**
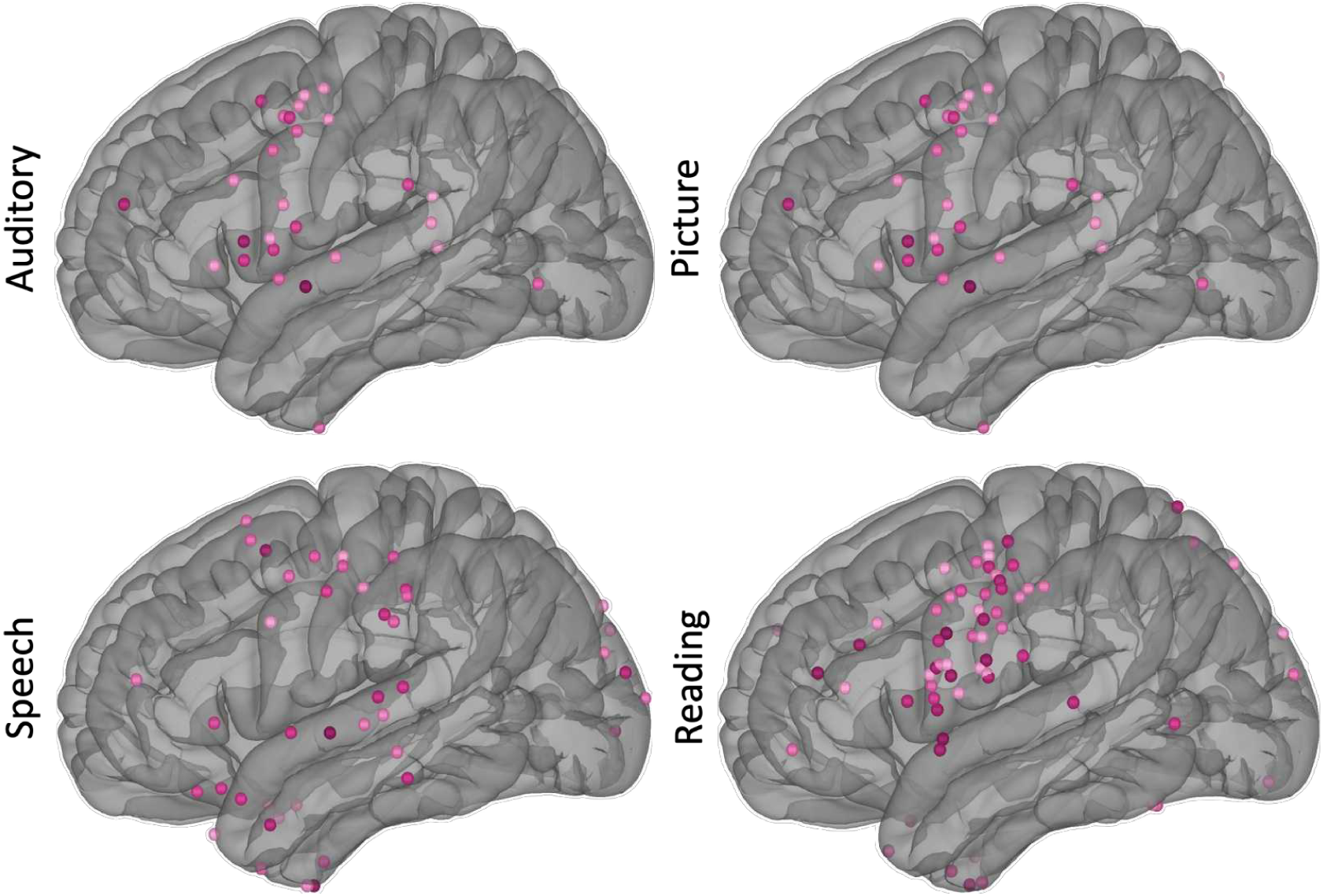
Distribution of electrodes exhibiting low high-gamma activity but high functional connectivity, identified as discriminative connections. These electrodes were selected using the relative distance index (RDI) and their color reflects the value of the RDI (see Figure 5F for the values before thresholding): all RDI values were pooled across states, and electrodes above the 90th percentile of this distribution were retained. Results are shown separately for each of the four states (Auditory, Picture, Speech, and Reading), projected onto a left-hemisphere lateral view.

**Supplementary Figure 9.**
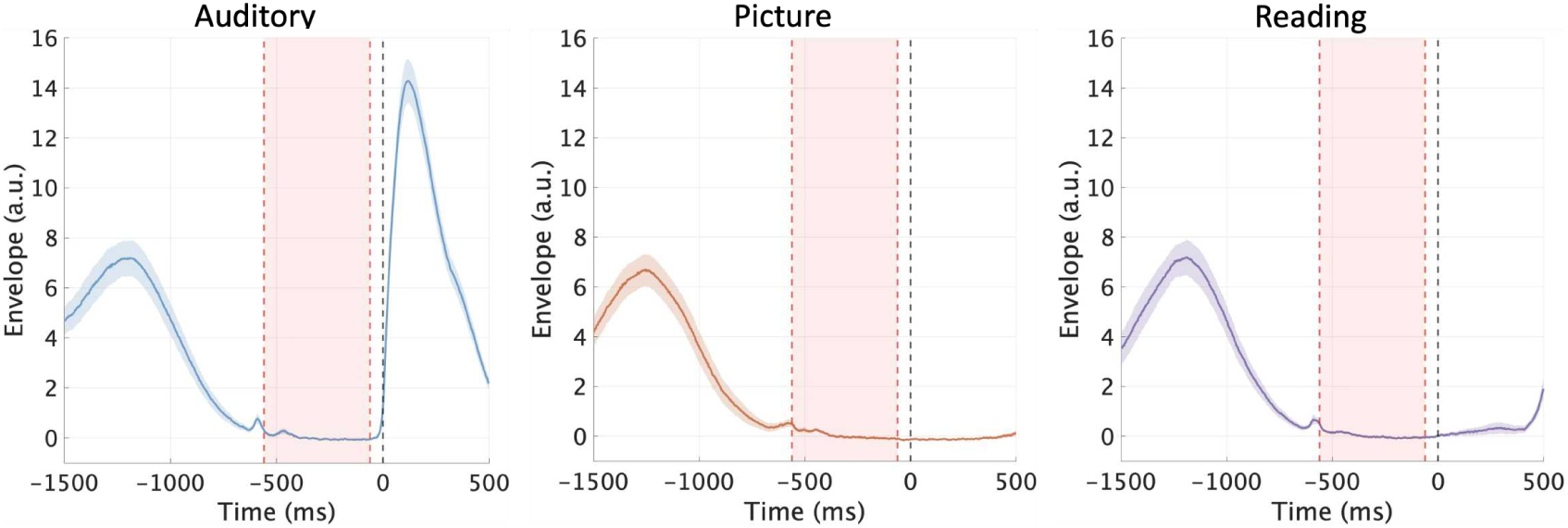
Average audio envelope across experimental tasks. Audio envelopes recorded during the auditory repetition (left), picture naming (middle), and visual word reading (right) tasks. Envelopes are epoched from −1500 ms prior to stimulus onset up to 500 ms post-onset, and averaged across trials and participants (shaded areas represent standard error). The small acoustic artifact visible around −600 ms corresponds to the experimenter pressing a keyboard button to advance the trial. This button press initiated a software random delay followed by a fixed 500 ms delay, ensuring a silent baseline window immediately preceding stimulus presentation. The shaded red area represents the selected baseline window in the analyses of this paper, starting from −560 ms to −60 ms prior to stimulus onset.

**Supplementary Table 2.** Electrode coverage by brain region. Number of analyzed recording sites in each region, summed across left- and right-hemisphere participants, together with the per-participant distribution. Counts reflect all the grid and strip electrodes before exclusion of bad electrodes, and for each participant only the hemisphere with the greater electrode count was retained, so every participant contributes to a single hemisphere. N is the number of participants with at least one electrode in the region; Median and Min is taken over those participants. The final row gives the total left- and right-hemisphere channel counts and the per-participant total-electrode median and range across all N = 38 participants (24 left-hemisphere, 14 right-hemisphere).

| Region | Number of participants | Left | Right | Total | Median | Range |
| --- | --- | --- | --- | --- | --- | --- |
| SuperiorFrontal | 38 | 427 | 290 | 717 | 17.5 | 3–43 |
| InferiorFrontal | 37 | 391 | 139 | 530 | 12 | 2–53 |
| precentral | 38 | 245 | 137 | 382 | 9 | 4–21 |
| postcentral | 38 | 287 | 133 | 420 | 11 | 6–28 |
| Parietal | 38 | 330 | 244 | 574 | 14.5 | 2–38 |
| SuperiorTemporal | 37 | 311 | 115 | 426 | 11 | 1–27 |
| InferiorTemporal | 38 | 422 | 208 | 630 | 18 | 1–34 |
| fusiform | 30 | 55 | 30 | 85 | 3 | 1–6 |
| Occipital | 20 | 53 | 29 | 82 | 4 | 1–10 |
| entorhinal | 25 | 32 | 18 | 50 | 2 | 1–5 |
| temporalpole | 13 | 15 | 6 | 21 | 1 | 1–3 |
| medialInferiorFrontal | 15 | 13 | 5 | 18 | 1 | 1–2 |
| parahippocampal | 10 | 8 | 5 | 13 | 1 | 1–2 |
| Cingulate | 2 | 3 | 0 | 3 | 1.5 | 1–2 |
| paracentral | 2 | 1 | 1 | 2 | 1 | 1–1 |
| Total | 38 | 2593 | 1360 | 3953 | 100 | 75–168 |

